# Human Breast Milk EVs Mitigate Endothelial Dysfunction: Preliminary Study

**DOI:** 10.1101/2024.05.20.594769

**Authors:** Young-Eun Cho, Shaoshuai Chen, Keith Crouch, Damon Shutt, Justin Kaufman, Brajesh Singh

## Abstract

**Background:** Endothelial cell (EC) dysfunction is an early indicator of failing vascular integrity, leading to various cardiovascular diseases. Toll-like receptor 4 (TLR4) activation is a key mechanism. Milk-derived extracellular vesicles (EVs) are known for their anti-inflammatory properties, particularly in suppressing TLR4 activation in damaged intestinal epithelial cells. This study explores the therapeutic potential of human breast milk EVs (bEVs) in EC dysfunction related to cardiovascular diseases.

**Methods:** Human breast milk EVs (bEVs) were isolated from healthy nursing mothers using ultracentrifugation. bEVs were applied to LPS-treated HUVECs, and the expression of inflammatory markers was measured using qPCR and western blotting. Angiogenesis was assessed via a wound assay. Additionally, bEVs were orally administered weekly for six weeks to high-fat diet-induced obese mice and lean mice. Metabolic phenotype characteristics and EC-dependent vasorelaxation were evaluated.

**Results:** bEV pre-treatment inhibited LPS-induced expression of inflammatory genes, including IL-6, IL-1b and VCAM-1. It also suppressed phospho-NFkB and TLR4 protein expression. bEVs enhanced EC migration, significantly increasing wound closure. Oral administration of bEVs restored impaired EC-dependent vasorelaxation in the mesenteric artery of obese mice, though metabolic parameters remained unchanged.

**Conclusion:** Our findings demonstrate the beneficial effects of bEVs on EC dysfunction, highlighting their potential as novel therapeutics for cardiovascular diseases. Future studies will focus on identifying specific bEV cargos and further evaluating their therapeutic effects on EC dysfunction.

## INTRODUCTION

One person dies every 33 seconds in the United States from cardiovascular diseases (CVDs)^1^. Endothelial cells (ECs) play a crucial role in maintaining cardiovascular homeostasis by controlling vascular reactivity, angiogenesis, permeability, blood clotting and more. [ref], Changes in EC function mark the onset of early CVD stages, consequently contributing to the progression of diverse cardiovascular conditions such as atherosclerosis, hypertension, coronary artery disease^2^. Increased inflammation is a hallmark of altered EC function. It contributes to increased oxidative stress and reduced vessel relaxation. It also contributes to impaired therapeutic angiogenesis in ECs, preventing the stimulation of new blood vessel growth to improve blood flow in ischemic tissues. Toll-like receptor 4 (TLR4) signaling plays a critical role in impairing these EC functions. TLR4 is highly expressed in microvascular ECs, and activated TLR4 is involved in promoting inflammatory response, leading to the development and progression of cardiovascular diseases^3-5^. Therefore, inhibiting TLR4 is considered a therapeutic strategy for CVD treatment.

Milk contains various components, such as proteins, peptides, lipids, and minerals, that have been shown to exhibit anti-inflammatory properties. Milk-derived bioactive peptides, milk proteins, and amino acids demonstrated reducing inflammatory gene expressions in ECs and EC-monocyte interactions via PPAR-γ dependent regulation of NF-κB ^6,7^. Milk also promotes angiogenesis [ref]. Growth factors, including vascular endothelial growth factor and epidermal growth factor, angiogenin-2, lactoferrin, and cadherin in milk, play a critical role in angiogenesis and proliferation ^8,9^[ref]. Furthermore, recently, milk has been identified as a rich source of extracellular vesicles (EVs). EVs are nanosized particles (30-150nm in diameter) containing various biological materials, including miRNAs, mRNAs, and DNAs^10,11^. Due to the nature of milk, milk EVs possess immune-promoting and anti-inflammatory properties, as evidenced in intestinal diseases like ulcerative colitis and necrotizing enterocolitis^12-14^. Several in vitro and in vivo studies demonstrated that milk EVs significantly reduced inflammation and apoptosis in intestinal epithelial cells, and recovered damaged intestinal structure^12,13,15^. This therapeutic effect was associated with TLR4 inhibition. Treatment with milk EVs led to a decrease in TLR4 expression, resulting in reduced levels of inflammatory markers such as NF-kB and NLRP3 inflammasome, as well as oxidative stress. These findings indicate that milk EVs hold significant therapeutic promise for alleviating EC inflammation and enhancing EC function. In this study, we explored the beneficial impacts of milk EVs on EC dysfunction using human breast milk. Our results provide insights into the potential benefits of milk EVs for breastfed infants and contribute foundational knowledge to the development of new therapeutics for cardiovascular diseases linked to EC dysfunction using milk EVs.

## Methods

### Breast milk EV collection

We recruited healthy nursing mothers who were ≥18 years old and had delivered a full-term (37–40 weeks) singleton newborn within the prior six months. Mothers with chronic conditions that might affect body weight changes, such as chronic diabetes or thyroid diseases, were excluded. After electronic informed consent was obtained, a breast milk sample collection kit was sent to the participant’s home with detailed instructions. After one hour of fasting, participants collected 50 ml of breast milk and froze it in their home freezer. Within 48 hours of sample collection, samples were returned to the lab and stored at -80°C until use. All procedures were followed under an approved IRB protocol (IRB #202005237). To avoid potential effects caused by the postpartum period and maternal BMI, milk from mothers whose pre-pregnancy BMI was either <25 (normal weight) and a postpartum day less than 60 days was used in this experiment. The demographics of donors of milk are described in Supplementary Table 1.

### Breast milk extracellular vesicle isolation and characterization

Breast milk EVs (bEVs) were isolated using ultracentrifugation described previously.^16^ Briefly, after removing fat and cell debris from the breast milk, the supernatant was centrifuged at 100,000 g for 2 hours to pellet EVs. The EV pellet was resuspended in sterile PBS and rotated overnight at 4°C. The concentration of bEV was measured with Qubit™ Protein and Protein Broad Range (BR) Assay Kits (Thermo Fisher Scientific, Waltham, MA, USA). The expression level of EV markers including CD9, CD81 and CD63 were measured using NanoView R100 and TSG101 was measured using a western blot.^16^

### HUVECs culture and bEV treatment

Human umbilical vein endothelial cells (HUVEC) were purchased from ATCC (Manassas, VA, USA) and cultured in EGM-2 Endothelial Cell Growth Medium-2 BulletKit (Lonza, Basel, Switzerland) at 37°C with 5% CO_2_ and 95% air conditions until passage 5. HUVECs were seeded on the tissue culture plates and incubated overnight. Then, bEVs or PBS were treated for 24 hours, followed by LPS 10μg/ml treatment for another 24 hours. Cells were harvested for the further processing of Western blotting or RT-qPCR.

### Cell viability assay

HUVECs were cultured for 12 hours at 96-well plates, then the cells were treated with different concentrations of bEVs (0 to 400 μg/ml) for 48 hours. Then, 15 μl of methyl thiazolyl tetrazolium (MTT, Promega, Madison, WI, USA.) was added to each well and incubated for another 4 hours. After removing the media, 150 μl of DMSO was added. The absorbance at 570 nm was assessed by a microplate reader.

### Quantitative real-time polymerase chain reaction

The RNA of HUVECs was isolated by miRVana kit (Thermo Fisher Scientific, Waltham, MA, USA). Then, the complementary DNA (cDNA) was synthesized by Prime Script RT Master Mix (Thermo Fisher Scientific), and a Taqman Master mix kit (Thermo Fisher Scientific) was used to perform quantitative real-time polymerase chain reaction (qRT-PCR) on the 7500 Real Time System (Applied Biosystems, CA, USA). GAPDH was employed as the internal controls. PCR primer sequences for each molecule are described in Supplementary Table 2. The 2^−ΔΔCt^ method was used to calculate the relative expression. The primers were synthesized by Integrated DNA Technologies (Coralville, IA, USA).

### EV internalization

To assess bEV internalization, bEVs were labeled with the fluorescent dye Dil according to the manufacturer’s instructions. Dil-labeled EVs were then incubated with HUVECs at a concentration of 50μg/ml for 24 hours. After incubation, the cells were washed three times with PBS to remove unbound EVs. Internalization of the exosomes was analyzed using fluorescence microscopy.

### Western blotting

Proteins were extracted from HUVECs using a RIPA buffer. Protein samples (25 μg) were subjected to SDS-PAGE, transferred on a PVDF membrane and then probed with primary antibodies; phospho-NF-κB p65 (Ser536, Cell Signaling), total NF-κB p65 (D14E12, Cell Signaling), TLR4 (25, SantaCruz), and β-actin (C4, SantaCruz). Protein level was visualized with enhanced chemiluminescence. The intensity of the bands was analyzed using Image J using a total NF-κB and β-actin as a control.

### Wound healing assay

To evaluate the beneficial effects of bEVs on the migration ability of HUVECs, a wound healing assay was performed. A straight scratch was introduced in the confluent cultures of HUVECs. After washing with PBS 3 times, cells were treated with bEVs 50μg/ml or vehicle controls. Time-lapse microscopy analysis for 48 hrs was performed with pictures taken every 15 min. The capacity of migration was evaluated by comparing the time taken and the percentage of wound closure area.

### Breast milk oral treatment

C57BL/6J male mice were purchased from The Jackson Laboratory (Bar Harbor, Maine, USA). Four-week old mice were randomly assigned to either high fat diet (HFD, 60% calories as fat, Envigo, Indianapolis, IN, USA) or normal chow diet (NCD, 10% calories as fat) groups. Food intake and body weight were measured weekly. For all experiments, mice were maintained on a normal 12:12-h light-dark cycle and provided regular mouse chow and water ad libitum at an Association for Assessment and Accreditation of Laboratory Animal Care (AAALAC)-accredited specific pathogen-free facility. After 14 weeks of diet, bEVs were orally administered to HFD-fed mice and NCD-fed mice weekly for 6 weeks via gavage (3ug of EVs/g). As a control, the same amount of PBS was administered to both HFD and NCD-fed mice.

### Metabolic function study

After 18 weeks of HFD and NCD, metabolic dysfunction was confirmed before and after bEV administration. The glucose tolerance test (GTT) was performed after overnight fasting (approximately 14 hours of fasting). Then glucose (2mg of 10% of glucose per body mass) was injected intraperitoneally, then blood glucose was measured from the tail-tip using a Care Touch glucometer (Future Diagnostics, USA) before, and at 0, 15, 30, 60, and 120 minutes after glucose administration. The insulin tolerance test (ITT) was performed after 8 hours of fasting. Insulin (0.75 IU insulin per body mass) was injected intraperitoneally. (Novolin R Insulin 100UN/ml, Novo Nordisk Inc., NJ) Then, blood glucose levels were measured from the tail-tip (kg) as described above. After completion of the test, the mice were returned to their home cage and given free access to food and water. Body fat composition was also measured before and after bEV treatment using a Nuclear Magnetic Resonance (LF50, Billerica, MA, USA).

### Myograph study

At the end of the 6-week treatment, EC-dependent vasorelaxation of mesenteric arteries was measured using a wire myograph (620M, DMT, Essen, Germany). Mesenteric arteries were dissected from the mice, and ring fragments were mounted in Krebs-Henseleit buffer (118.4 mM NaCl, 4.7 mM KCl, 2.5 mM CaCl2, 1.2 mM KH2PO4, 1.2 mM MgSO4, 25 mM NaHCO3, 11.1 mM glucose). The organ bath was maintained at 37°C, and the pH was adjusted using a gas mixture of 5% CO_2_ and 95% O_2_ throughout the experiment. Each ring was initially stretched to an optimal load of 2.0 mN. After a 30-minute equilibration period, the viability of the arteries was tested with 70 mM KCl. This was followed by treatment with 10^−4^ M phenylephrine. Then, serial treatments with acetylcholine (ACh, 10^−9^ M to 10^−4^ M) were then applied to examine EC-dependent vasorelaxation. At the end of the experiment, the arteries were treated with 10^−4^ M sodium nitroprusside (SNP) to assess smooth muscle function. The percentage of relaxation was compared between groups.

### Statistical analysis

Demographics and experimental measures between groups were compared by the Mann-Whitney U test with a statistical threshold of *p* = 0.05 using SPSS 27 (IBM, NY, USA) or GraphPad Prism 7 (GraphPad Software, Inc., La Jolla, CA, USA). All plots were generated using GraphPad Prism 7 (GraphPad Software, Inc.). p<0.05 was considered to indicate a statistically significant difference.

## RESULTS

Prior to examining the effect of bEVs on HUVECs, the effect of bEVs on cell viability was tested. We tested 5 different bEV samples with 6 different concentrations. As shown in Fig 1, the survival rate of HUVECs was not altered by bEV pre-treatment at concentrations of 0, 25, and 50 μg/ml. However, the concentration of bEVs at 100, 200 and 400 μg/ml reduces the cell viability. Therefore, we determined the use 50 μg/ml of bEVs for the anti-inflammatory effect in HUVECs.

**Fig 1.**
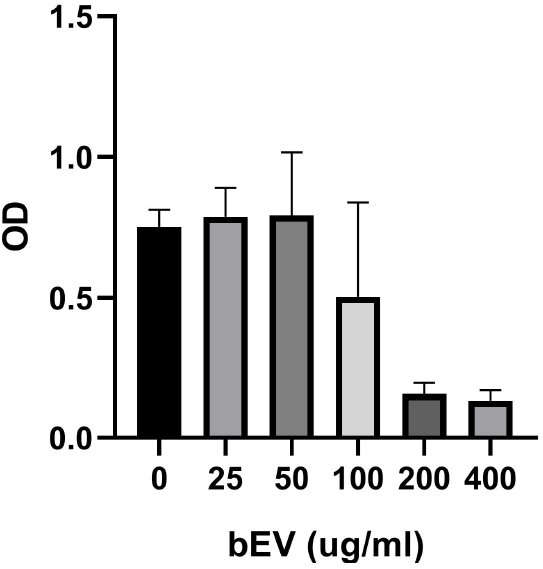
Assessment of Cell Viability of HUVECs with bEVs. MTT assay was performed with different concentrations of bEVs. Based on the result, we determined to test 50μg/ml of bEVs to treat HUVECs.

Before we tested anti-inflammatory effect of bEVs on HUVECs, first, we confirmed the internalization of bEVs in HUVECs (Fig 2). Then, the expression of TLR-4 downstream pro-inflammatory molecules including IL-6, IL-1b, and VCAM-1 was measured using qPCR following treatment with 10 μg/ml LPS in the presence or absence of bEVs. We tested bEV samples (n=8-10). The gene expression was significantly reduced with the pre-treatment of bEVs (Fig 3A-C). The phospho-NFkB level and TLR4 expression increased by LPS treatment, which also decreased with bEV treatment (Fig 3D-E).

**Fig 2.**
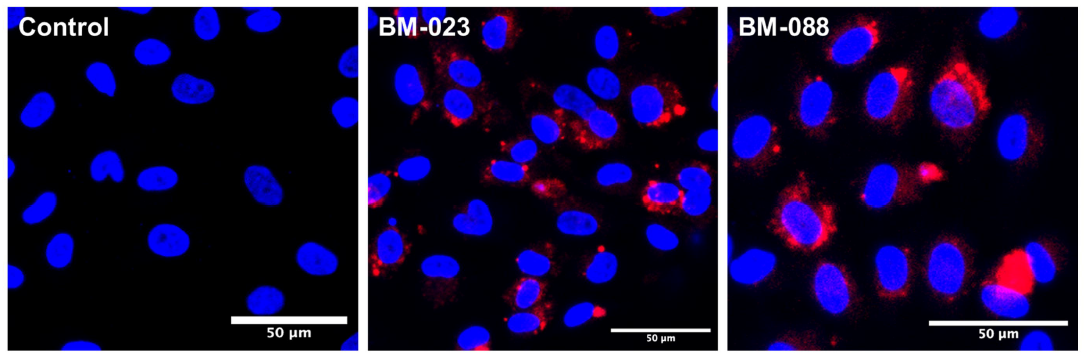
Internalization of bEVs to HUVECs. bEV internalization was visualized in HUVECs. Cells were treated with 50μg/ml of Oil-labelled bEVs (red). After 24 hrs, cells were fixed and stained for nuclei (blue).

**Fig 3.**
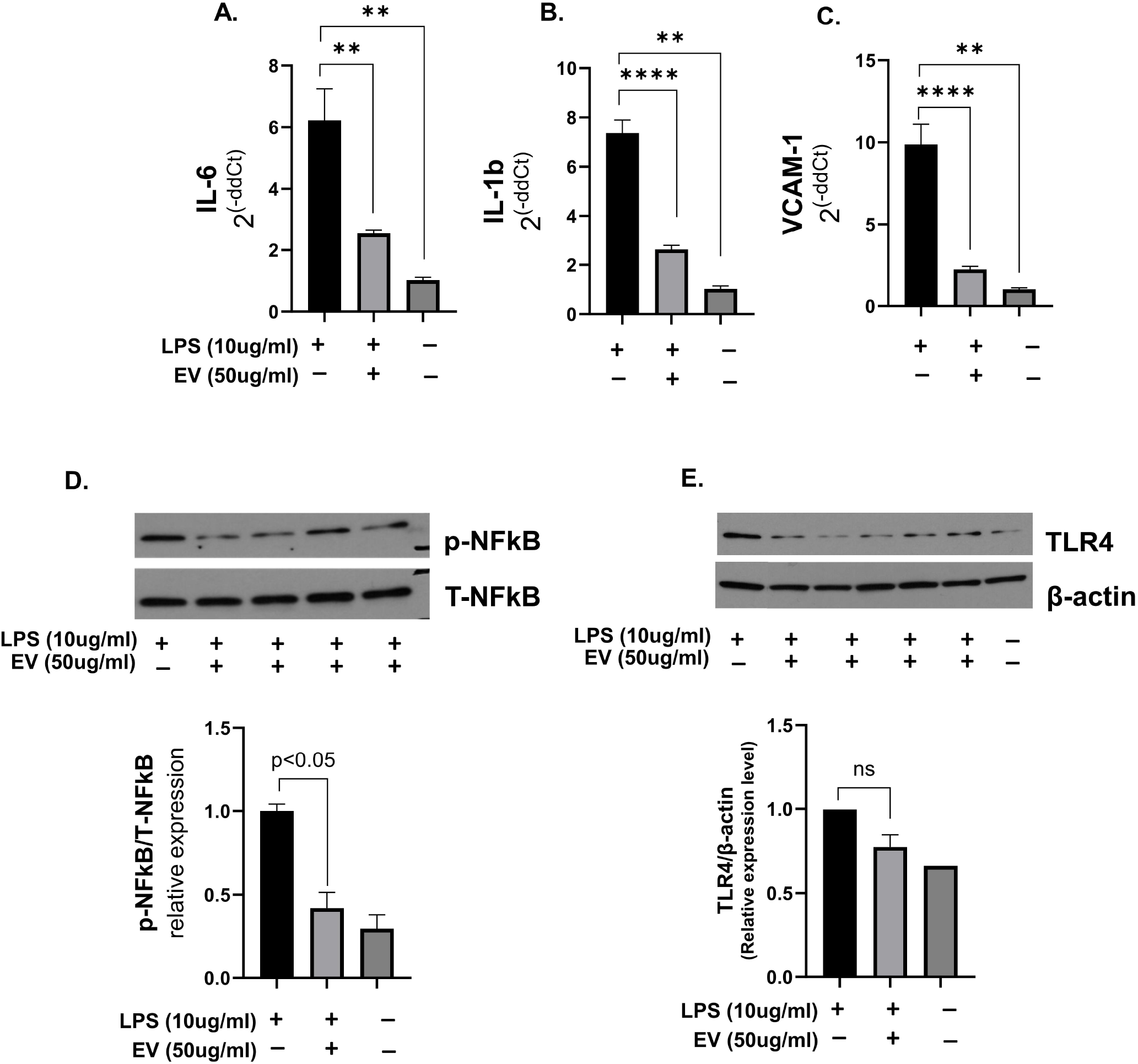
Reduced inflammatory gene expression by bEV pretreatment in LPS-treated HUVECs. Inflammatory gene expression was measured using qPCR **(A-C)**. Also, phospho-NFkB and TLR4 protein expression levels were measured using western blotting **(D and E)**.

bEVs also improved the migration ability of ECs (Fig 4). bEVs-treated HUVECs demonstrated significantly faster wound closure, compared to PBS-treated HUVECs, suggesting that bEVs may have a potential role in enhancing angiogenesis in ECs.

**Fig 4.**
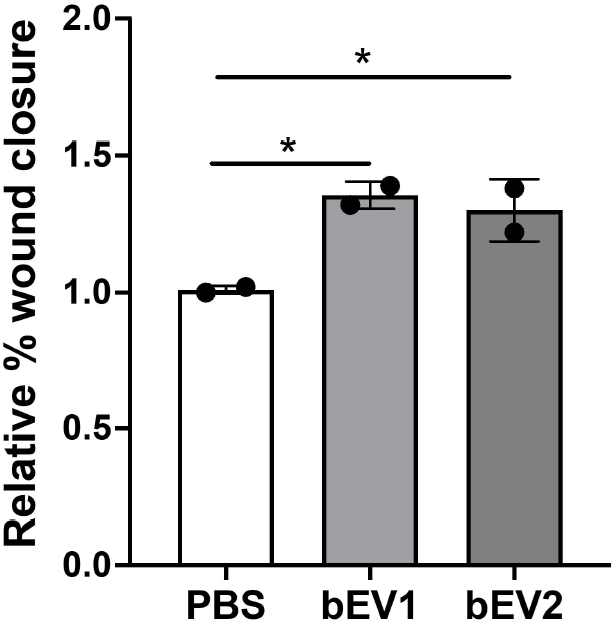
Improved angiogenesis in bEV-treated HUVECs. A wound healing assay showed the effects of bEVs on HUVECs migration analyzed as relative % wound closure (n = 2 in duplicate). Confluent cultures of HUVECs were treated with bEVs (n=2). *p < 0.05

Finally, we test the effect of bEVs on high-fat diet-induced obese mice. bEVs were administered orally weekly for six weeks, then metabolic parameters including body weight, glucose tolerance, insulin tolerance and body fat composition, were examined before and after bEV treatment. The parameter did not show any differences (Suppl Fig 1) after bEV treatment. Then, endothelial-dependent vasorelaxation was examined from the mesenteric artery on the wire myograph. PBS-treated obese mice showed impaired ACh-dependent vasorelaxation; however, it was restored in obese mice who were bEV-administered (Fig 5).

**Fig 5.**
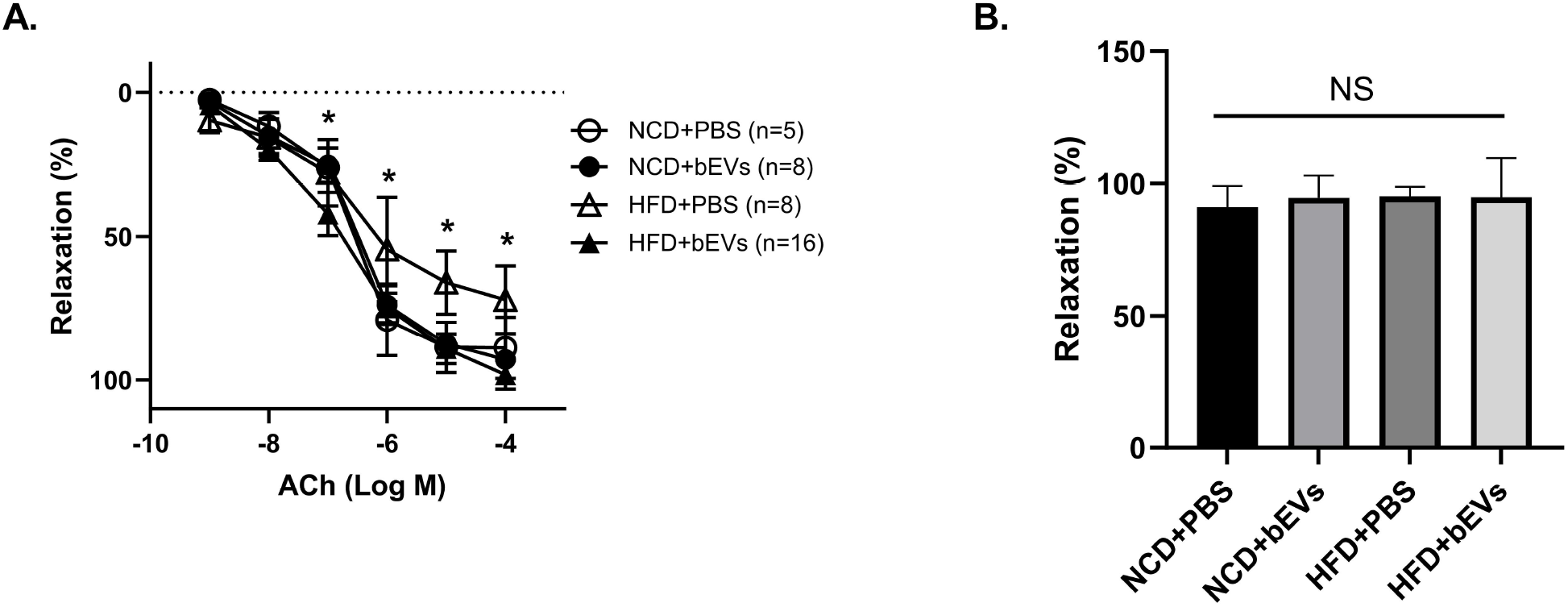
Improved EC-dependent vasorelaxation in bEV-treated obese mice. High-fat diet-induced obese mice (HFD mice) and normal chow diet mice (NCO mice) orally received bEVs (3μg/g) weekly for 6 weeks. Then ACh-induced EC-dependent vasorelaxation was evaluated from mesenteric arteries (ACh 10^9^M-10^−4^M). The EC-dependent vasorelaxation, which was significantly impaired in obese mice, was recovered in bEV treatment mice **(A)**. However, SNP (10^−4^M) induced vasorelaxation was not changed **(B)**. N indicates the number of experiments. * indicates p<0.05 between HFD mice fed PBS and HFD mice fed bEVs. NS indicates no significance.

## DISCUSSION

Milk EVs are known to have anti-inflammatory properties as a part of the milk component. They effectively reduced the inflammatory gene expression in intestinal epithelial cells, however, it has not been tested in the other types of cells. In this study, using human breast milk EVs, we demonstrated the beneficial role of milk EVs on EC dysfunction, which is a critical phenomenon in many kinds of CVDs. To our knowledge, it is the first study demonstrating direct evidence showing the beneficial effect of bEVs in ECs.

We confirmed that the anti-inflammatory effect of bEVs is valid in ECs. LPS treatment can mimic the chronic inflammatory condition in obese individuals^17^, which are commonly linked to EC dysfunction and cardiovascular diseases. LPS treatment increased IL-6 and IL-1b gene expression, TLR4 downstream signaling pro-inflammatory molecules, which decreased when bEVs were pre-treated. IL-6 is a multipotent cytokine and increases the proliferation and differentiation of numerous cell types. In ECs, IL-6 is involved in the inflammatory process, which results in atherosclerotic plaque development and plaque destabilization^18^. IL-1b also facilitates early atherogenic lesion formation by increasing monocyte adhesion to ECs via upregulation of adhesion molecules, such as VCAM-1^19,20^. Both IL-6 and IL-1b also directly affect NOS activity and expression in ECs, leading to inactivation of NOS and NO bioavailability. Increased p-NFkB leads to increased inflammatory gene expression, which was also reduced by bEV treatment. It is correlated with the in vivo findings obtained from the obese mice, restoring EC-dependent relaxation after oral administration of bEVs. These findings are also congruent with previous studies testing bovine milk EVs on intestinal epithelial cells^12,15^.

While milk is known for its potent angiogenic effects, studies have primarily focused on testing milk EVs as carriers for therapeutic drugs in wound healing. Only a few studies have directly examined the impact of milk itself in this context. A study evaluating the angiogenic and wound-healing potential of bovine milk EVs showed results similar to those observed in our study^21,22^. Kim et al. compared the angiogenic and wound-healing potency of colostrum and mature milk from cows, finding that colostrum outperformed mature milk in promoting re-epithelialization, activating angiogenesis, and enhancing extracellular matrix maturation^21^. However, tube formation and cell proliferation assays yielded similar results for EVs derived from both types of milk. Zhang et al. also tested bovine milk EVs on cardiac fibrosis^22^. Milk EVs alleviated cardiac fibrosis and enhanced cardiac function by enhancing angiogenesis. O Other studies have examined engineered milk EVs aimed at augmenting their wound-healing effectiveness, either by integrating a hydrogel material or by encapsulating potential therapeutics such as miRs^23-25^. However, the comprehensive content of milk EVs remains incompletely understood, highlighting the necessity for additional research to harness milk EVs for therapeutic purposes or as carriers for therapeutics.

Another way to develop novel therapeutics from bEVs would be to identify the key molecules responsible for this beneficial effect. miRNAs are one of the major cargo molecules of bEVs. We recently profiled the miRNAs of bEVs obtained from 65 healthy nursing mothers^16^. Of 798 miRNAs screened, we found that miR-30b-5p and miR-494-3p were highly expressed in bEVs, suggesting they might be potential candidates for this beneficial effect on ECs. miR-30b-5p is suggested as a key immune-regulatory miRNA of milk EVs^26^. miR-30b-5p is implicated in cardiovascular and metabolic diseases by modulating inflammation and oxidative stress^27-29^. Notably, miR-30b-5p inhibits TLR4 activity, decreasing the expression of pro-inflammatory molecules including VCAM-1, ICAM-1, TNF-α, NFκB, and p38 in ECs ^29,30^. Previous studies have demonstrated that miR-494-3p has multifaceted therapeutic potentials across various pathological conditions, including inflammatory responses, oxidative stress pathways, and angiogenesis^31-33^. Interestingly, in WI-38 cells, miR-494 directly interacts with TLR4 and regulates oxidative stress^31^. This evidence supports the potential of these miRNAs as future novel therapeutics. Future studies investigating their roles in ECs will be needed for the development of novel therapeutics.

We were not able to observe the changes in metabolic parameters of obese mice after the oral administration of bEVs. It is well-known that breastfeeding in infants was associated with a reduced risk of obesity^34^. The beneficial effect of breast milk on infant health and development may be associated with various factors, including its nutrient composition, hormonal factors such as leptin, and microbiome changes^35^. Since breast milk contains a significant amount of EVs, bEVs may play an important role in various effects on infants; however, this relationship is not yet fully understood. Currently, there is no direct evidence showing that milk EVs contribute to a reduced risk of obesity or alter the characteristics of adipocytes or tissues. Studies demonstrate that adipogenesis-regulating miRNAs, such as let-7a and miR-378, are found in milk^36^, and suggest that milk EV miRNAs may alter gene expression in the infant’s gut to regulate their metabolism and growth^37^. In our pilot study, we administered one dose of bEVs weekly for 6 weeks, but we did not observe any changes in metabolic phenotypes. It is possible that the dosage or duration of administration was not sufficient to induce metabolic phenotype changes. In vitro studies using 3T3-L1 cells or primary cultured adipocytes from obese mice, with varying doses of bEVs treatment, as well as animal studies involving different doses with long-term administration, are needed to elucidate the role of bEVs in metabolic parameters. This research will provide valuable insight into understanding the role of bEVs in breastfed infants.

In conclusion, our study has shown that bEVs possess a strong anti-inflammatory effect and an angiogenic effect in ECs. Further research is needed to identify the key cargo molecules of bEVs responsible for this beneficial effect. This knowledge will facilitate the development of novel therapeutics using EVs as a precise and effective tool for targeted treatments of cardiovascular diseases associated with impaired ECs.

## REFERENCES

1 NIDDK. Overweight & Obesity Statistics.

2 Kotsis, V., Stabouli, S., Papakatsika, S., Rizos, Z. & Parati, G. Mechanisms of obesity-induced hypertension. Hypertens Res 33, 386–393 (2010). 10.1038/hr.2010.9

3 Kawasaki, T. & Kawai, T. Toll-like receptor signaling pathways. Front Immunol 5, 461 (2014). 10.3389/fimmu.2014.00461

4 Nunes, K. P., de Oliveira, A. A., Lima, V. V. & Webb, R. C. Toll-Like Receptor 4 and Blood Pressure: Lessons From Animal Studies. Front Physiol 10, 655 (2019). 10.3389/fphys.2019.00655

5 Hernanz, R. et al. Toll-like receptor 4 contributes to vascular remodelling and endothelial dysfunction in angiotensin II-induced hypertension. Br J Pharmacol 172, 3159–3176 (2015). 10.1111/bph.13117

6 Marcone, S., Haughton, K., Simpson, P. J., Belton, O. & Fitzgerald, D. J. Milk-derived bioactive peptides inhibit human endothelial-monocyte interactions via PPAR-gamma dependent regulation of NF-kappaB. J Inflamm (Lond) 12, 1 (2015). 10.1186/s12950-014-0044-1

7 R., M. S. C. B. O. B. I. Milk Proteins and Amino Acids Modulate Inflammatory Gene Expression in Vascular Endothelial Cells. The FASEB Journal 30, 1169 (2016). 10.1096/fasebj.30.1_supplement.1169.10

8 Medina, M. A. & Quesada, A. R. Dietary proteins and angiogenesis. Nutrients 6, 371–381 (2014). 10.3390/nu6010371

9 Somasundaram, I., Kaingade, P.; Bhonde, R. Ch. Breast Milk Critical Secretary Growth Factors for Angiogenesis, Cell Proliferation and Tissue Homeostasis, (Springer, Singapore, 2023).

10 Gurung, S., Perocheau, D., Touramanidou, L. & Baruteau, J. The exosome journey: from biogenesis to uptake and intracellular signalling. Cell Commun Signal 19, 47 (2021). 10.1186/s12964-021-00730-1

11 Elliott, R. O. & He, M. Unlocking the Power of Exosomes for Crossing Biological Barriers in Drug Delivery. Pharmaceutics 13 (2021). 10.3390/pharmaceutics13010122

12 Tong, L. et al. Milk-derived extracellular vesicles alleviate ulcerative colitis by regulating the gut immunity and reshaping the gut microbiota. Theranostics 11, 8570–8586 (2021). 10.7150/thno.62046

13 Xie, M. Y. et al. Porcine Milk Exosome MiRNAs Attenuate LPS-Induced Apoptosis through Inhibiting TLR4/NF-kappaB and p53 Pathways in Intestinal Epithelial Cells. J Agric Food Chem 67, 9477–9491 (2019). 10.1021/acs.jafc.9b02925

14 Gao, H. N. et al. Yak milk-derived exosomes alleviate lipopolysaccharide-induced intestinal inflammation by inhibiting PI3K/AKT/C3 pathway activation. J Dairy Sci 104, 8411–8424 (2021). 10.3168/jds.2021-20175

15 Good, M. et al. Breast milk protects against the development of necrotizing enterocolitis through inhibition of Toll-like receptor 4 in the intestinal epithelium via activation of the epidermal growth factor receptor. Mucosal Immunol 8, 1166–1179 (2015). 10.1038/mi.2015.30

16 Cho, Y. E. et al. Extracellular vesicle miRNAs in breast milk of obese mothers. Front Nutr 9, 976886 (2022). 10.3389/fnut.2022.976886

17 Troseid, M. et al. Plasma lipopolysaccharide is closely associated with glycemic control and abdominal obesity: evidence from bariatric surgery. Diabetes Care 36, 3627–3632 (2013). 10.2337/dc13-0451

18 Su, J. H. et al. Interleukin-6: A Novel Target for Cardio-Cerebrovascular Diseases. Front Pharmacol 12, 745061 (2021). 10.3389/fphar.2021.745061

19 Gomez, D. et al. Interleukin-1beta has atheroprotective effects in advanced atherosclerotic lesions of mice. Nat Med 24, 1418–1429 (2018). 10.1038/s41591-018-0124-5

20 Kihara, T. et al. Interleukin-1beta enhances cell adhesion in human endothelial cells via microRNA-1914-5p suppression. Biochem Biophys Rep 27, 101046 (2021). 10.1016/j.bbrep.2021.101046

21 Kim, H. et al. Harnessing the Natural Healing Power of Colostrum: Bovine Milk-Derived Extracellular Vesicles from Colostrum Facilitating the Transition from Inflammation to Tissue Regeneration for Accelerating Cutaneous Wound Healing. Adv Healthc Mater 11, e2102027 (2022). 10.1002/adhm.202102027

22 Zhang, C. et al. Bovine Milk Exosomes Alleviate Cardiac Fibrosis via Enhancing Angiogenesis In Vivo and In Vitro. J Cardiovasc Transl Res 15, 560–570 (2022). 10.1007/s12265-021-10174-0

23 Fan, L. et al. Antioxidant-Engineered Milk-Derived Extracellular Vesicles for Accelerating Wound Healing via Regulation of the PI3K-AKT Signaling Pathway. Adv Healthc Mater 12, e2301865 (2023). 10.1002/adhm.202301865

24 Yan, C. et al. Milk exosomes-mediated miR-31-5p delivery accelerates diabetic wound healing through promoting angiogenesis. Drug Deliv 29, 214–228 (2022). 10.1080/10717544.2021.2023699

25 Xiang, X. et al. Milk-derived exosomes carrying siRNA-KEAP1 promote diabetic wound healing by improving oxidative stress. Drug Deliv Transl Res 13, 2286–2296 (2023). 10.1007/s13346-023-01306-x

26 Zhou, Q. et al. Immune-related microRNAs are abundant in breast milk exosomes. Int J Biol Sci 8, 118–123 (2012). 10.7150/ijbs.8.118

27 Yang, L. et al. Cannabinoid Receptor 1/miR-30b-5p Axis Governs Macrophage NLRP3 Expression and Inflammasome Activation in Liver Inflammatory Disease. Mol Ther Nucleic Acids 20, 725–738 (2020). 10.1016/j.omtn.2020.04.010

28 Sun, Y. et al. Regulatory Role of rno-miR-30b-5p in IL-10 and Toll-like Receptor 4 Expressions of T Lymphocytes in Experimental Autoimmune Uveitis In Vitro. Mediators Inflamm 2018, 2574067 (2018). 10.1155/2018/2574067

29 Demolli, S. et al. MicroRNA-30 mediates anti-inflammatory effects of shear stress and KLF2 via repression of angiopoietin 2. J Mol Cell Cardiol 88, 111–119 (2015). 10.1016/j.yjmcc.2015.10.009

30 Zhou, Z. et al. MicroRNA-30-3p Suppresses Inflammatory Factor-Induced Endothelial Cell Injury by Targeting TCF21. Mediators Inflamm 2019, 1342190 (2019). 10.1155/2019/1342190

31 Dai, Z., Hu, J., Luo, Z. & Xiao, J. Downregulation of circ_0035292 Alleviates LPS-Induced WI-38 Cell Injury via Targeting miR-494-3p/TLR4 Pathway in Infantile Pneumonia. Biochem Genet (2023). 10.1007/s10528-023-10455-0

32 Wang, H., Wang, S. & Huang, S. MiR-494-3p alleviates acute lung injury through regulating NLRP3 activation by targeting CMPK2. Biochem Cell Biol 99, 286–295 (2021). 10.1139/bcb-2020-0243

33 Liu, H. et al. Dendritic cell-derived exosomal miR-494-3p promotes angiogenesis following myocardial infarction. Int J Mol Med 47, 315–325 (2021). 10.3892/ijmm.2020.4776

34 Abbas, M. A., Al-Saigh, N. N. & Saqallah, F. G. Regulation of adipogenesis by exosomal milk miRNA. Rev Endocr Metab Disord 24, 297–316 (2023). 10.1007/s11154-023-09788-3

35 Mantzorou, M. et al. Exclusive Breastfeeding for at Least Four Months Is Associated with a Lower Prevalence of Overweight and Obesity in Mothers and Their Children after 2–5 Years from Delivery. Nutrients 14 (2022). 10.3390/nu14173599

36 Xi, Y. et al. The levels of human milk microRNAs and their association with maternal weight characteristics. Eur J Clin Nutr 70, 445–449 (2016). 10.1038/ejcn.2015.168

37 Shah, K. B. et al. Human Milk Exosomal MicroRNA: Associations with Maternal Overweight/Obesity and Infant Body Composition at 1 Month of Life. Nutrients 13 (2021). 10.3390/nu13041091

