## Supplementary figures and images for "Human Breast Milk EVs Mitigate Endothelial Dysfunction: Preliminary Study"

### Supple Fig1

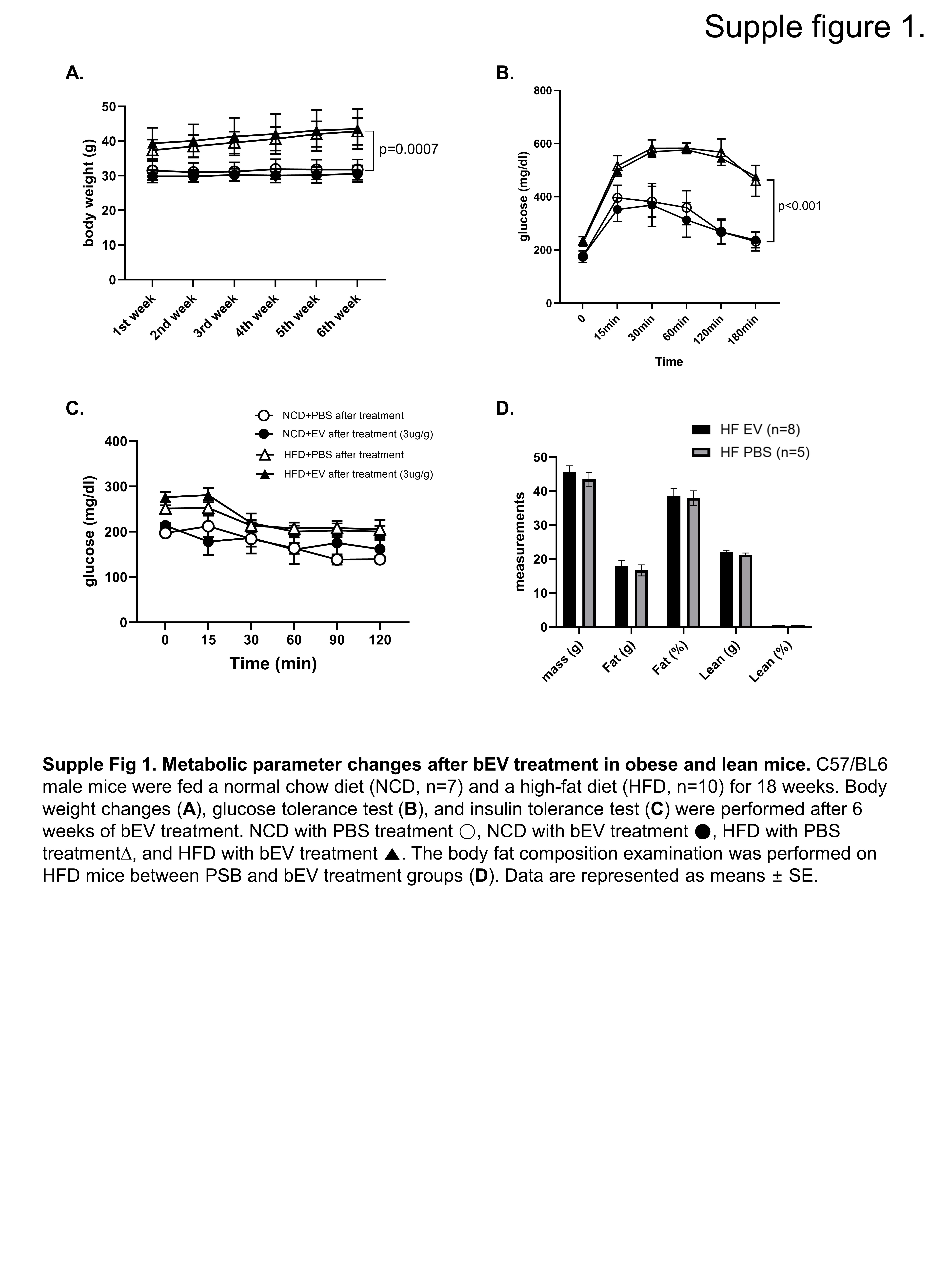

### Supple Table 1

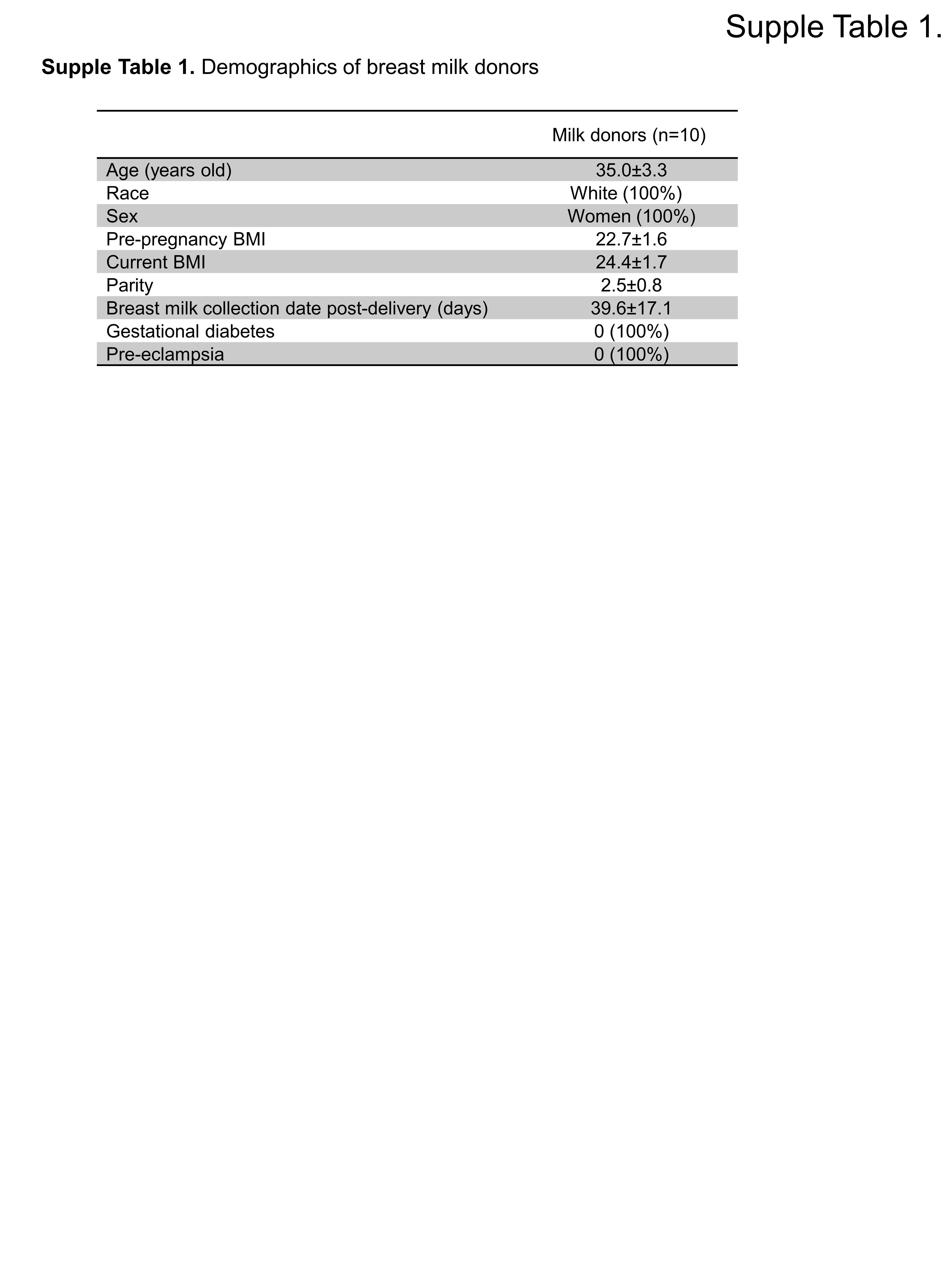

### Supple Table 2

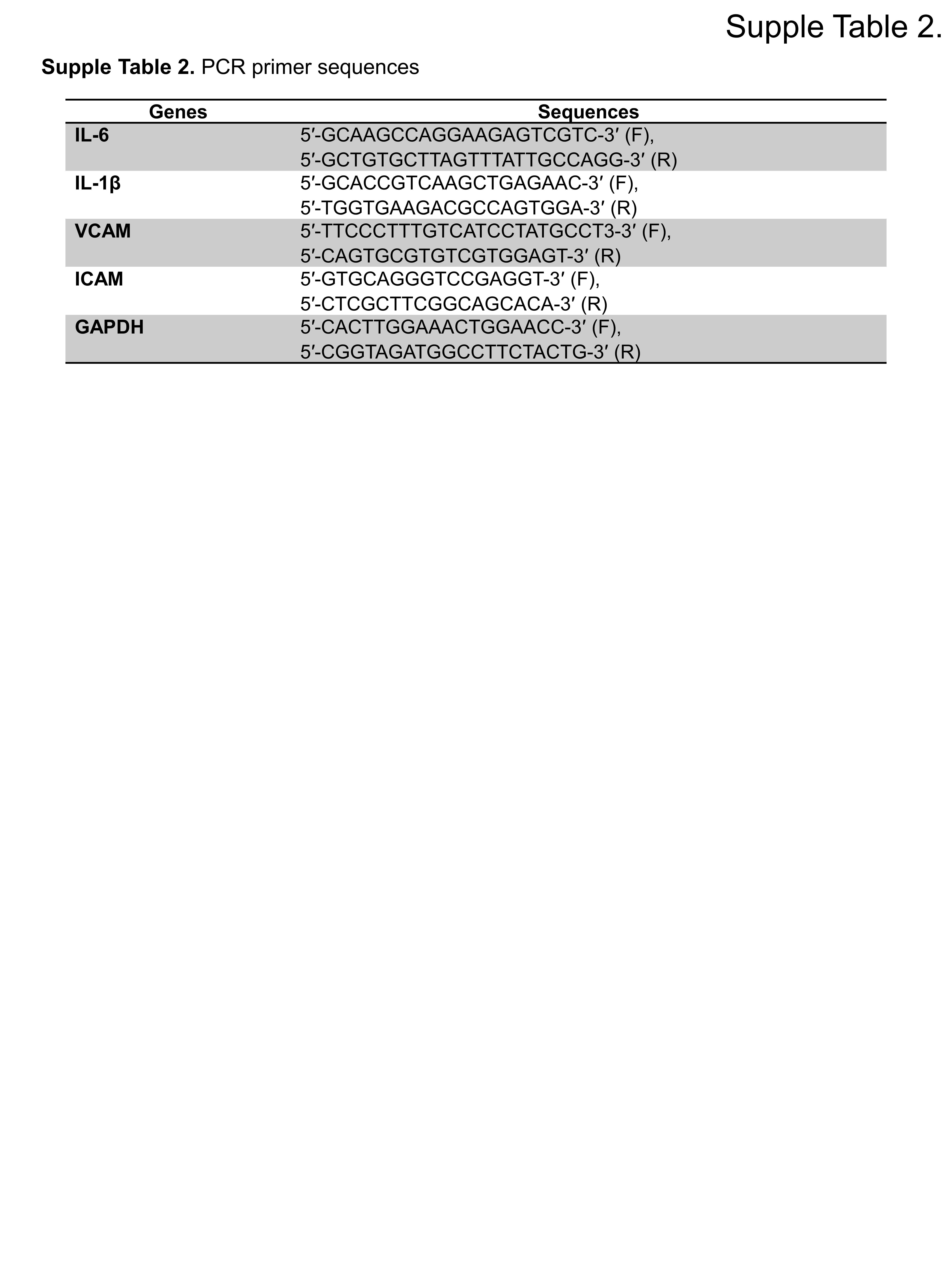
